# Convergence in amino acid outsourcing between animals and predatory bacteria

**DOI:** 10.1101/2025.02.22.639622

**Authors:** Niko Kasalo, Mirjana Domazet-Lošo, Tomislav Domazet-Lošo

**Author notes:** Correspondence: T.D.-L.

## Abstract

All animals have outsourced about half of the 20 proteinogenic amino acids (AAs). We recently demonstrated that the loss of biosynthetic pathways for these outsourced AAs is driven by energy-saving selection. Paradoxically, these metabolic simplifications enabled animals to use costly AAs more frequently in their proteomes, allowing them to explore sequence space more freely. Based on these findings, we proposed that environmental AA availability and cellular respiration mode are the two primary factors determining the evolution of AA auxotrophies in animals. Remarkably, our recent analysis showed that bacterial AA auxotrophies are also governed by energy-related selection, thereby roughly converging with animals. However, bacterial AA auxotrophies are highly heterogeneous and scattered across the bacterial phylogeny, making direct ecological and physiological comparisons with the animal AA outsourcing model challenging. To better test the universality of our model, we focused on Bdellovibrionota and Myxococcota—two closely related bacterial phyla that, through aerobic respiration and a predatory lifestyle, best parallel animals. Here, we show that Bdellovibrionota, driven by energy-related selection, outsourced a highly similar set of AAs to those in animals. This sharply contrasts with Myxococcota, which exhibit far fewer AA auxotrophies and rarely show signatures of energy-driven selection. These differences are also reflected in Bdellovibrionota proteomes, which are substantially more expensive than those of Myxococcota. Finally, we found evidence that the expression of costly proteins plays a crucial role in the predatory phase of the Bdellovibrio life cycle. Together, our findings suggest that Bdellovibrionota, through their obligate predatory lifestyle, exhibit the closest analogy to the AA auxotrophy phenotype observed in animals. In contrast, facultative predation, as seen in Myxococcota, appears to substantially limit the evolution of AA auxotrophies. These cross-domain convergences strongly support the general validity of our AA outsourcing model.

## 1. Introduction

The biosynthesis of amino acids (AAs) is a fundamental biochemical process essential for sustaining all life on Earth. However, despite the universal metabolic demand for all 20 proteinogenic AAs, some organisms have lost the ability to synthesize certain AAs endogenously. The most well-known example is animals [1–3], which have lost the capacity to produce approximately half of the complete AA set. A similar pattern of auxotrophies has independently emerged in other eukaryotic groups, such as certain amoebae and euglenozoans [1,2]. While most bacteria remain fully prototrophic [4], many bacterial lineages display varying degrees of AA auxotrophy [4–6].

It has long been speculated that the ability to acquire AAs from the environment influences the evolution of AA auxotrophies [2]. For instance, experiments have shown that bacteria auxotrophic for an externally supplemented AA can gain a selective advantage over fully prototrophic counterparts [7]. However, a robust theoretical framework for this phenomenon remained elusive. In our previous work, we addressed this gap by proposing a model that explains the evolution of AA outsourcing through several key factors: an AA is more likely to be lost if (i) its biosynthesis is highly energy-demanding, (ii) it has a low pleiotropic effect, (iii) it is abundantly available in the environment, and (iv) the organism relies on efficient aerobic respiration for energy production [1].

Most importantly, we demonstrated that there is a constant selective pressure to outsource the synthesis of energetically costly AAs to the environment [1]. Surprisingly, this leads to an increased usage of these expensive AAs in proteomes, allowing animal proteins to explore sequence space more freely [1,8]. To determine whether these global patterns are also valid in bacteria, we investigated AA auxotrophies across bacterial phylogeny and found that energy-related selection also plays an important role in shaping AA outsourcing in bacteria [5]. However, bacteria exhibit far greater metabolic and ecological diversity than animals [4,9,10], which necessitates testing our AA outsourcing model in specific bacterial groups whose lifestyles more closely parallel those of animals.

One such group is the phylum Bdellovibrionota, which includes several aerobic or microaerophilic species with an obligate predatory lifestyle [11–13]. Within this phylum, predation occurs via two distinct strategies: epibiotic predation, where the bacterium attaches to the prey cell and leeches nutrients from its surface, and endobiotic predation, where the bacterium enters the prey cell and forms a bdelloplast, within which it feeds, grows, and divides [11–13]. The life cycle of Bdellovibrionota consists of two transcriptionally distinct phases: the attack phase, during which the bacterium seeks and attaches to prey, and the feeding phase, characterized by growth and replication [12,14].

Another bacterial group exhibiting animal-like behaviors is the phylum Myxococcota, a close relative of Bdellovibrionota [15,16]. Myxococcota are aerobes known for their complex multicellular behaviors, including social movement in coordinated “wolf packs,” fruiting body formation, and predation [17]. However, unlike Bdellovibrionota, Myxococcota are not obligate predators—they can scavenge nutrients from dead organic matter and survive periods of starvation through sporulation [18–20].

The predatory lifestyle evolved independently in Bdellovibrionota and Myxococcota [16], leading to vastly different phenotypic outcomes [11–13,18–20]. Thus, these two bacterial groups provide an ideal system to test the predictive power of our AA outsourcing model, as they allow us to directly link the evolution of AA auxotrophies in bacteria to distinct ecological and physiological traits [1,5,8].

Here, we demonstrate that Bdellovibrionota evolved a remarkably animal-like set of AA auxotrophies, accompanied by an increase in relative proteome costs. Our findings indicate that energy-related selection played a key role in shaping these auxotrophies and that the attack and feeding phases of their life cycle exhibit distinct energy dynamics at the transcriptomic level. Surprisingly, this pattern does not hold for *Myxococcota*, which exhibit fewer auxotrophies and lower proteome costs. This suggests a fundamental difference in how diverse predatory lifestyles shape metabolic evolution in these two groups.

## 2. Results

### 2.1. AA Auxotrophies in Bdellovibrionota and Myxococcota

We suspected that the animal-like ecophysiology of Bdellovibrionota and Myxococcota imposes energy-related selection on their AA metabolism, leading to reductions in AA biosynthetic pathways similar to those observed in animals. To test this hypothesis, we first estimated the completeness of AA biosynthesis pathways in 89 Bdellovibrionota and 203 Myxococcota high-quality proteomes, which we retrieved from the NCBI database (Table S1). To assess AA biosynthesis pathway completeness, which represents the likelihood of a given pathway being present, we used the MMseqs2 clustering approach [5,21]. Heatmap representations clearly revealed significantly lower AA completeness scores (CS) in Bdellovibrionota compared to Myxococcota (Figure 1, File S1).

**Figure 1.**
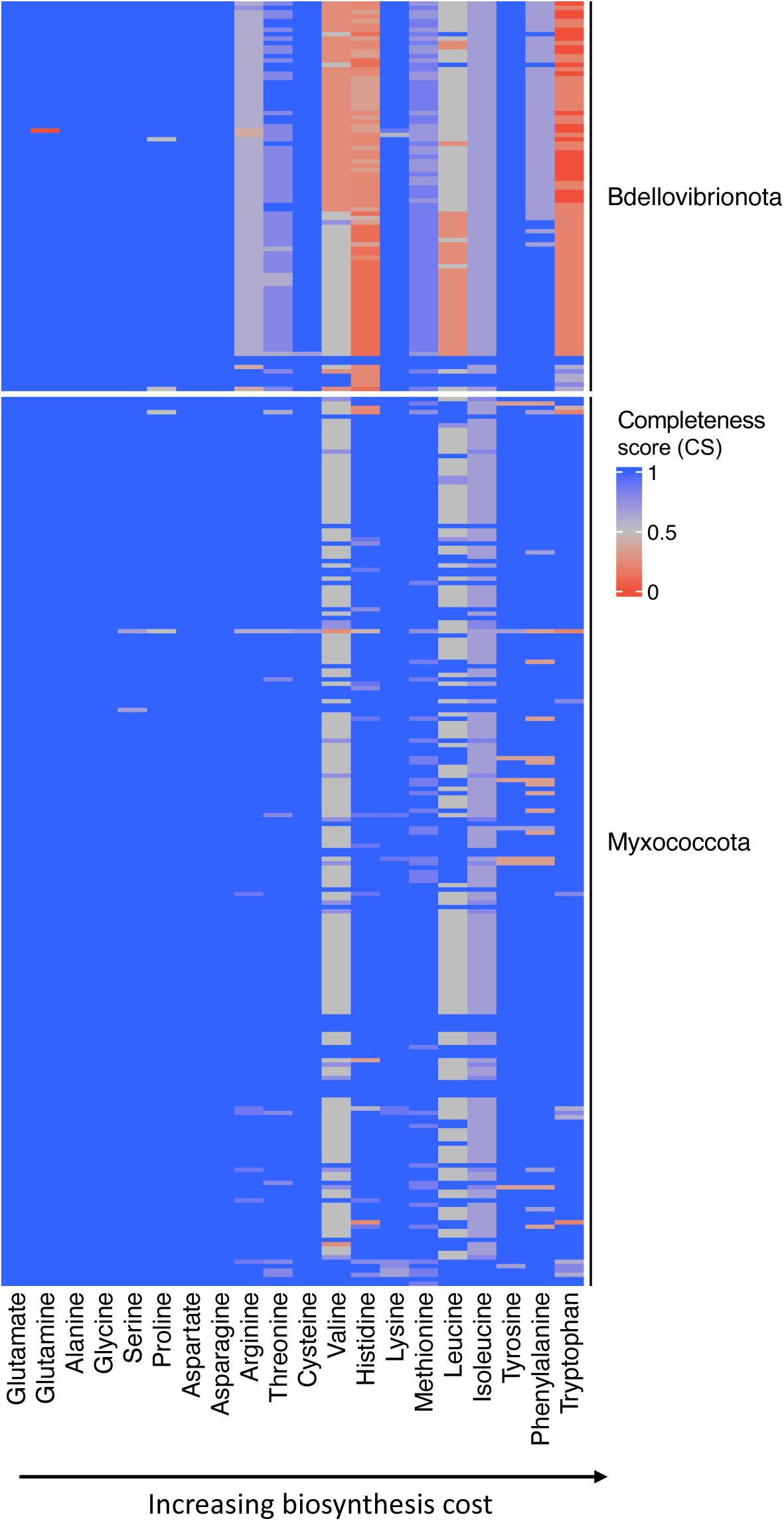
Completeness score of AA biosynthesis pathways in Bdellovibrionota and Myxococcota. We created a database of 89 Bdellovibrionota and 203 Myxococcota proteomes to get a comprehensive overview of AA dispensability in these groups. Full figure is shown in File S1. We retrieved all enzymes involved in AA biosynthesis from the KEGG and MetaCyc databases (reference collection) and searched for their homologs within our Bdellovibrionota/Myxococcota database using MMseqs2 (see Methods). For each AA, we showed a completeness score (CS), which represents the percentage of enzymes within a pathway that returned significant sequence similarity matches to our reference collection of AA biosynthesis enzymes. In the case of AAs with multiple alternative pathways, we showed the results only for the most complete one.

Most Bdellovibrionota showed reductions in pathway completeness scores for nine amino acids (AAs) that fall on the expensive end of the biosynthesis cost distribution (Figure 1, File S1). This set of AAs with low completeness scores largely overlaps with those that are auxotrophic in animals [1], with one notable exception—lysine, which appears to be prototrophic in Bdellovibrionota (Figure 1, File S1). In contrast, the pattern of AA biosynthesis pathway reduction in Myxococcota is markedly different. Most Myxococcota remain prototrophic for the majority of AAs, with only valine, leucine, and isoleucine biosynthesis pathways consistently exhibiting reductions (Figure 1, File S1). An exception to this trend is observed in two Myxococcota species—*Vulgatibacter incomptus* and *Pajaroellobacter abortibovis*—which display a substantial number of AA auxotrophies (Figure S1). Taken together, these findings suggest that while *Myxococcota* abolish the production of some expensive AAs, specific ecological factors likely prevent them from outsourcing most of their costly AA biosynthesis pathways.

### 2.2. Expensive AAs Are Commonly Outsourced in Bdellovibrionota

To globally test whether energy-related selection influences the observed reductions in AA biosynthesis pathways of Bdellovibrionota and Myxococcota, we correlated the average AA auxotrophy index with opportunity costs (OC) calculated under high respiration mode [5], a metric that estimates the impact of AA biosynthesis on the cell’s energy budget. To obtain the average AA auxotrophy index for a given AA, we first subtracted the completeness score from 1 (Material and Methods, Equation 1) and then averaged the resulting auxotrophy index (AI) values across all considered proteomes. We detected a significant correlation between higher biosynthesis costs and the loss of AA biosynthetic ability in Bdellovibrionota (Figure 2a). This suggests that selection driven by energy management shaped the global pattern of auxotrophies in Bdellovibrionota. As might be expected considering heatmap pattern (Figure 1), this broad-scale analysis did not detect a positive correlation in Myxococcota, which are prototrophic for most AAs (Figure 2b).

**Figure 2.**
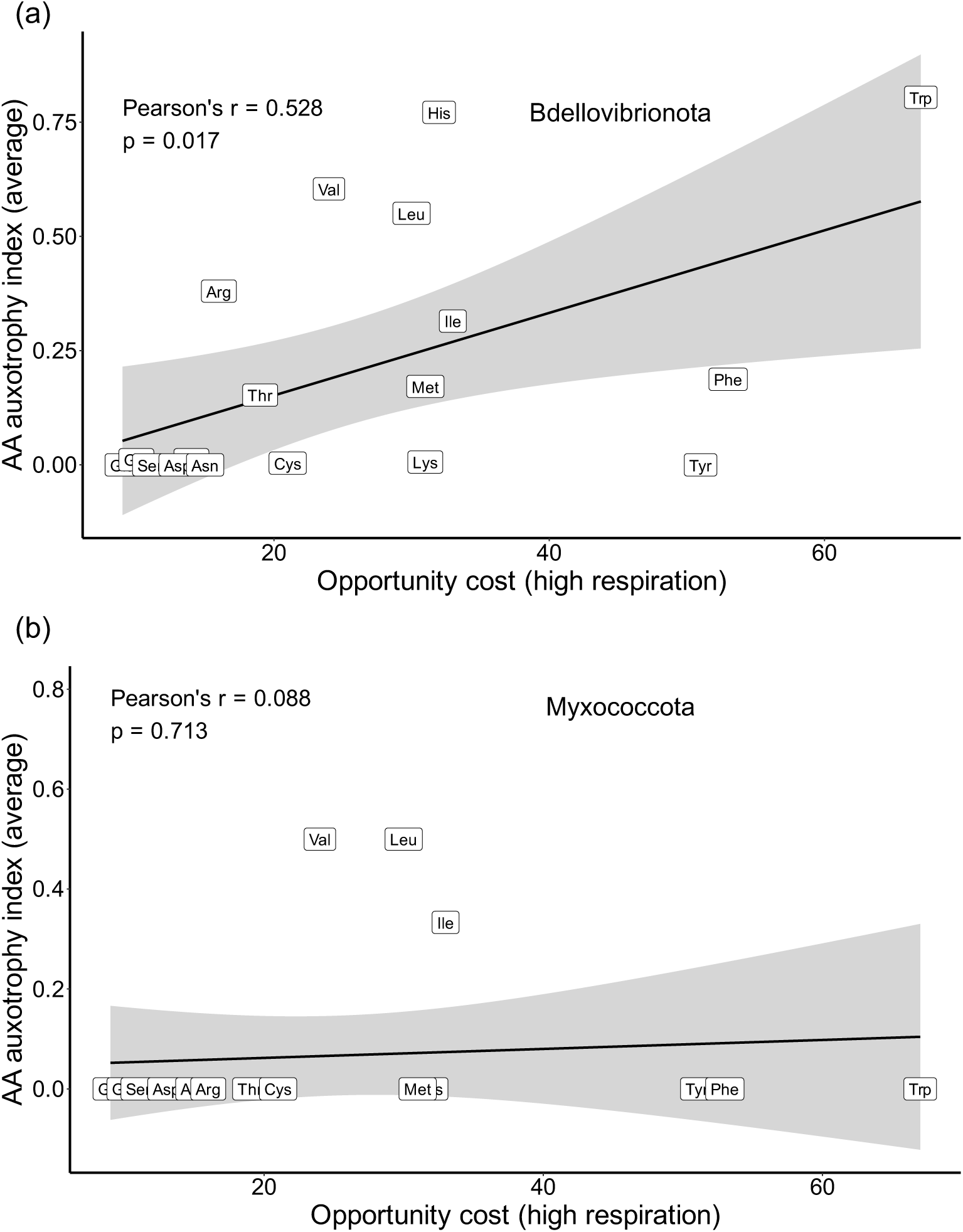
Correlation between AA biosynthesis cost and AA auxotrophy index. We estimated the AA auxotrophy index (AI), a measure which equals one minus completeness score (see Materials and Methods, Equation 1), for 89 Bdellovibrionota (**a**) and 203 Myxococcota (**b**) proteomes and calculated the average value for every AA. We correlated this value with the opportunity cost (OC) of each AA, calculated for high respiration mode (see Materials and Methods). Pearson correlation coefficient and p-value are shown on the graph.

### 2.3. Energy-optimizing selection drives AA auxotrophies in Bdellovibrionota

To further evaluate whether the observed reductions in AA biosynthesis pathways in Bdellovibrionota and Myxococcota result from energy-optimizing selection, we analyzed AA auxotrophy patterns at the species level. To achieve this, we first explicitly determined the AA auxotrophy status of each species by transforming the completeness score (CS) to binary values (auxotrophic/prototrophic) (see Materials and Methods, Table S2). Although this procedure reduces the information contained in the completeness scores, it allowed us to test the impact of energy-related selection more directly by explicitly defining auxotrophic and prototrophic AAs. In principle, the loss of even a single enzyme within a biochemical pathway could render that pathway nonfunctional. Thus, we assigned auxotrophy status to any AA whose biosynthesis pathway was incomplete. Using this transformed dataset, we statistically compared opportunity costs between auxotrophic and prototrophic AAs within each species [1]. In addition, we applied a permutation-based selection test to assess the probability that the observed constellation of auxotrophic AAs evolved under energy-related selection [1].

On average, Bdellovibrionota species exhibited 7.89 auxotrophic AAs per species (Table S2). The comparison between auxotrophic and prototrophic AA sets using the Mann-Whitney non-parametric test shows that auxotrophic AAs have significantly higher opportunity costs in 92% of the 89 tested Bdellovibrionota species (Table S2; File S2). For illustration, we singled out the results of this comparison for the type strains of *Bdellovibrio bacteriovorus* and *Pseudobdellovibrio exovorus*, representing endobiotic and epibiotic lifestyles, respectively (Figure 3). It is evident that, regardless of feeding ecology, auxotrophic AAs are significantly more expensive than prototrophic ones (Figure 3a,b). The permutation-based selection test revealed similar global trends, detecting that energy-related selection impacted the observed distribution of auxotrophic AAs in 61% of Bdellovibrionota species (Table S2; File S2). For instance, the average opportunity cost of the auxotrophic AA sets observed in *Bdellovibrio bacteriovorus* and *Pseudobdellovibrio exovorus* falls at the right end of the distribution of all possible permutations (Figure 3c,d). This indicates, just like in animals [1], that energy-optimizing selection governed the outsourcing of auxotrophic AAs in these bacteria.

**Figure 3.**
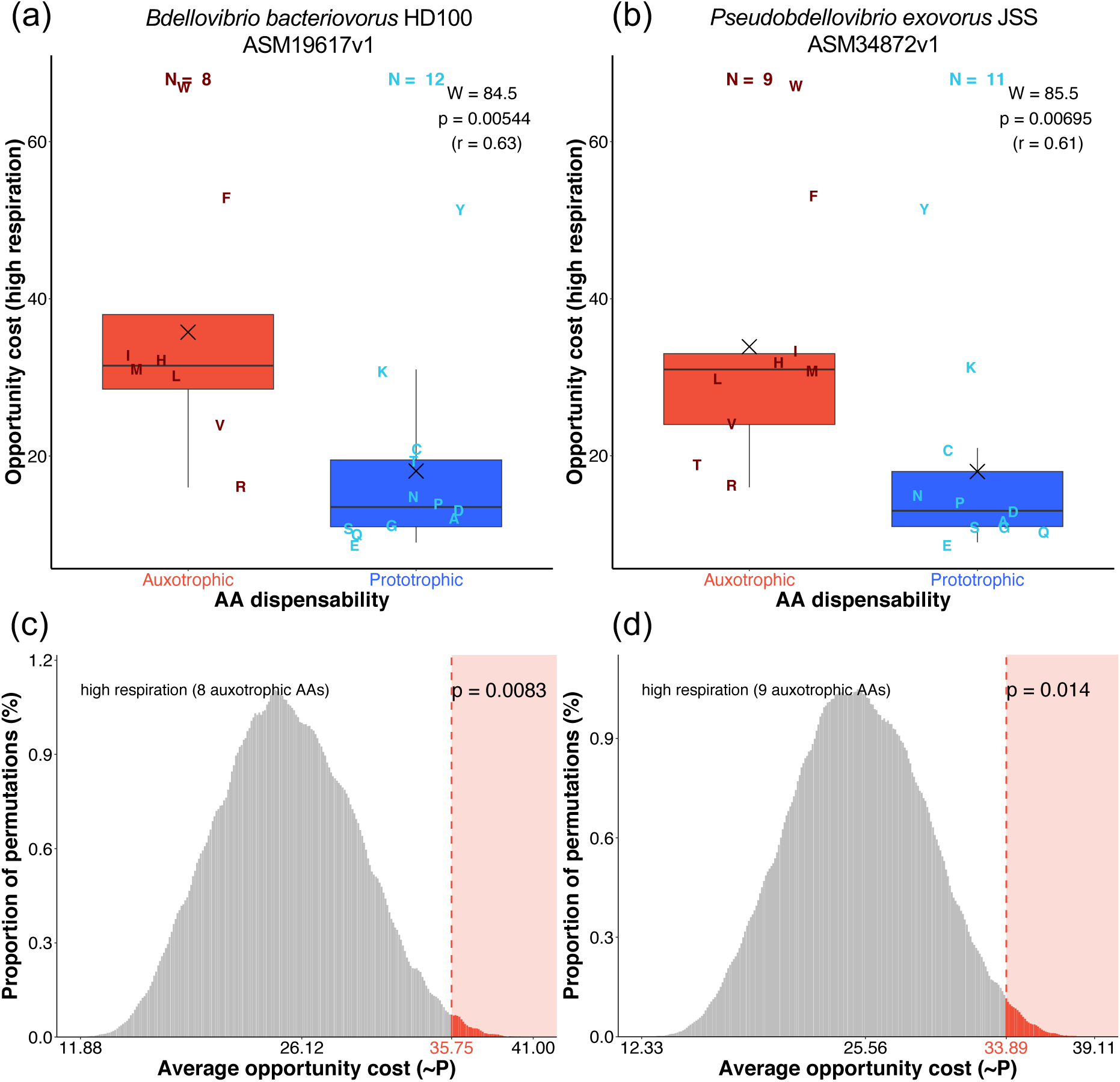
Comparisons of auxotrophic and prototrophic AA sets in *Bdellovibrio bacteriovorus* (a, c) and *Pseudobdellovibrio exovorus* (b, d). (a, b) The comparison of opportunity costs (OC) in high respiration mode between auxotrophic and prototrophic AA groups was tested by the Mann-Whitney U test with continuity correction. The corresponding W-value, p-value, and effect size (r) are depicted in each panel. The X symbol represents the mean. Individual AAs are shown by one-letter symbols. **(c, d)** Permutation analyses of opportunity costs (selection tests) [1] were conducted by calculating the average opportunity cost for every possible permutation within the number of auxotrophic AAs identified in each species (*n* = 8 for *B. bacteriovorus*, *n* = 9 for *P. exovorus*). The proportions of these averages are shown in histograms. The obtained distribution represents the empirical probability mass function (PMF). The value in red denotes the average opportunity cost (OC) value of the auxotrophic AA sets observed in nature. The p-value was calculated by summing the proportions of permutations with average opportunity cost values equal to or more extreme than the observed opportunity cost value. Low p-values indicate a high probability that energy-related selection drove the loss of auxotrophic AA biosynthesis capability.

The trends are quite different in Myxococcota, where an average of 2.79 AAs are auxotrophic (Table S2). The Mann-Whitney non-parametric test shows that auxotrophic AAs have significantly higher opportunity costs in only 9% of the 203 tested species (Table S2; File S2). In comparison, the permutation-based selection test revealed similar results, indicating that energy-related selection impacted AA auxotrophies in only 7% of Myxococcota species (Table S2). A prominent representative of Myxococcota that shows the impact of energy-related selection on AA auxotrophies is *Pajaroellobacter abortibovis*, a species whose pathogenic ecology differs from the facultative predatory lifestyle of most other Myxococcota (File S1, Table S2, File S2). Taken together, these findings suggest that energy-optimizing selection related to AA auxotrophies is rather rare among Myxococcota compared to Bdellovibrionota.

### 2.4. Bdellovibrionota Encode Expensive Proteomes

The fact that Bdellovibrionota converge to an animal-like set of amino acid (AA) auxotrophies, while closely related Myxococcota remain mainly prototrophic, allows us to further test the predictions of our model [1]. In our previous work, we demonstrated that animals encode significantly more expensive proteomes compared to choanoflagellates, their sister group, which is primarily prototrophic [1]. Based on this finding, we proposed that animals have costlier proteomes than choanoflagellates, likely because they expend less energy on amino acid biosynthesis, allowing them to maintain a larger number of expensive auxotrophic AAs in their proteomes [1]. Using a non-parametric test to compare the opportunity costs of an average AA in bacterial proteomes (OC*_proteome_*, Table A1, Equation 2), we recovered an analogous result: the proteomes of the more auxotrophic Bdellovibrionota are significantly more expensive than those of the more prototrophic Myxococcota (Figure 4), underscoring the universality of energy-related selection on AA composition in proteomes.

**Figure 4.**
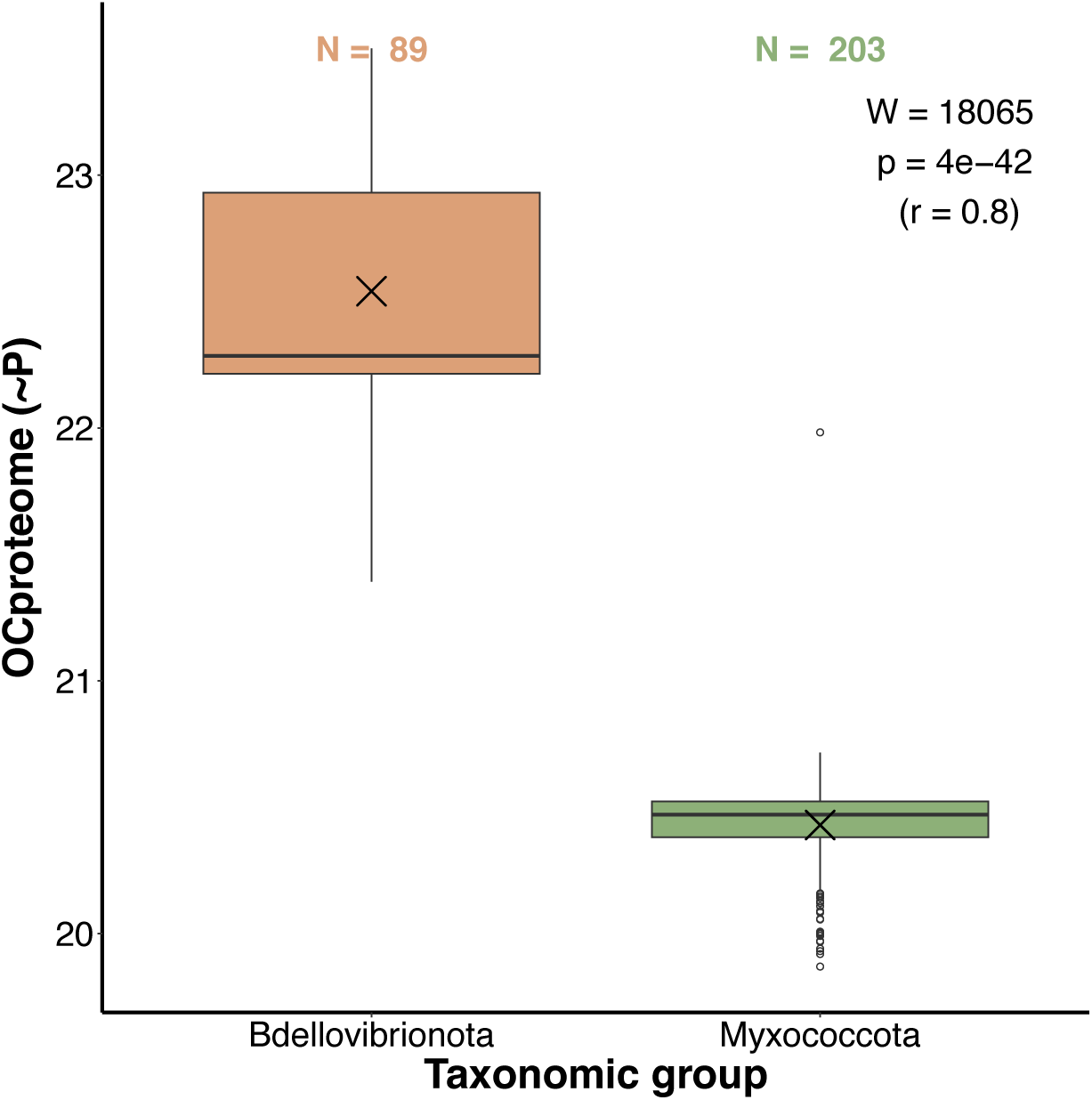
Comparison of the proteome opportunity cost (OC*_proteome_*) between Bdellovibrionota and Myxococcota. The opportunity cost under high respiration mode of an average AA in a proteome (OC*_proteom_*, Equation 2) represents a weighted mean of AA biosynthesis energy costs where the frequencies of twenty AAs in the proteome act as weights. The differences in energy costs between the two groups were shown by boxplots and the significance of these differences was tested by the Mann-Whitney U test with continuity correction. We depicted the corresponding W-value, p-value, and effect size (r) in each panel. The X symbol represents the mean. The list of Bdellovibrionota (89) and Myxococcota (203) proteomes used in the calculations is available in File S1.

### 2.5. Expensive Proteins in Bdellovibrio Drive Active Predation

We used the published transcriptome data to examine the energetics of the two distinct phases in the life cycle of *Bdellovibrio bacteriovorus*: the attack phase and the growth phase [14]. For each transcript, we calculated its frequency in the transcriptome at a given life cycle phase [22]. We then multiplied this transcript frequency by the opportunity cost of an average amino acid (AA) encoded by that transcript (OC*_protein_*) (Equations 3 and 4). This measure, which we named the transcript energy score (TES, Table A1), couples transcript levels in the cell with the encoded protein costs (see Material and Methods). Higher TES values reflect greater impact, while lower TES values indicate a smaller impact of a given transcript on the total energy budget of the cell. We performed the TES-based analysis in two ways: by including all genes expressed per phase and by considering only those which were exclusively expressed in one phase (Figure 5).

**Figure 5.**
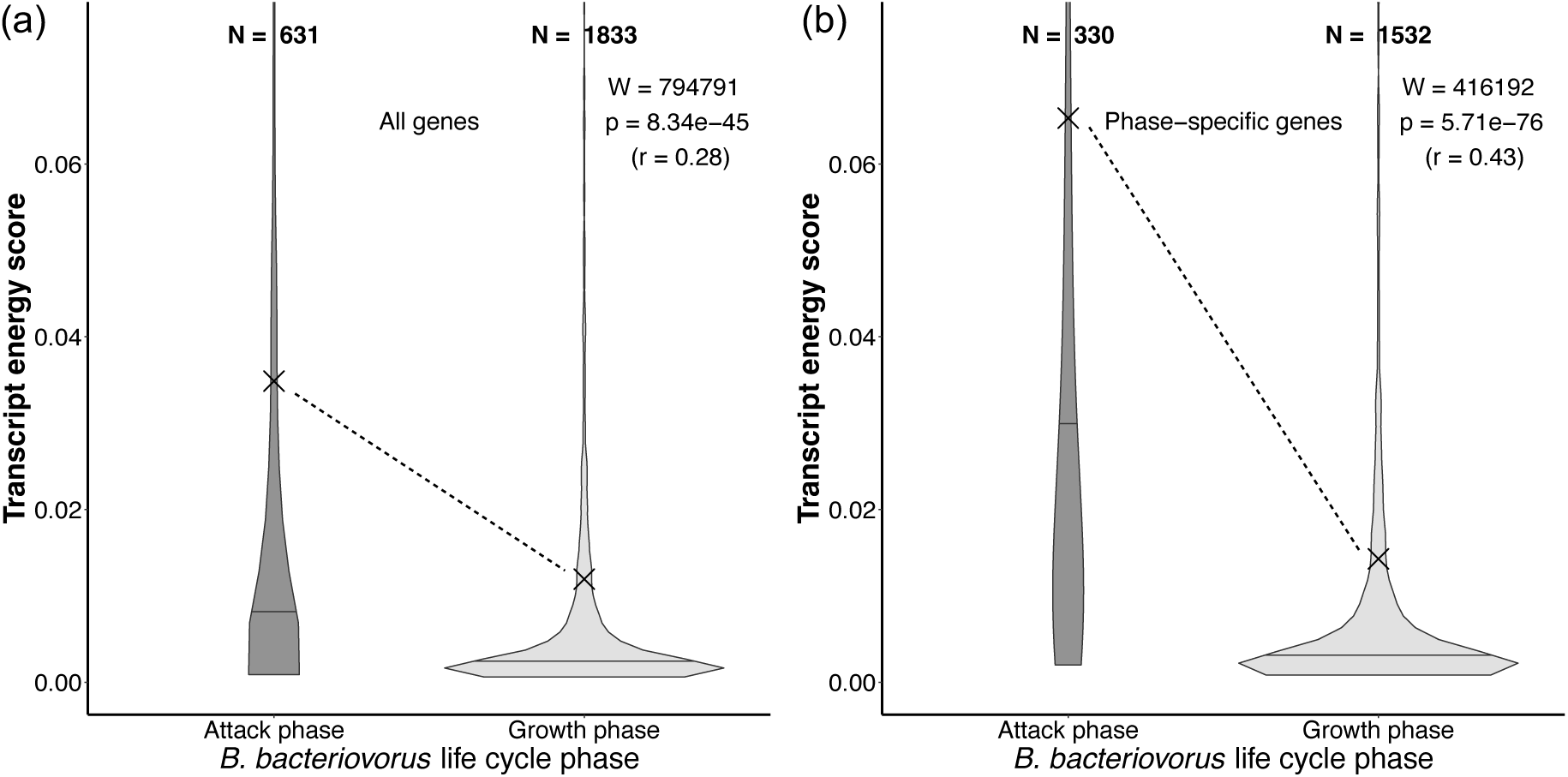
The comparison of transcriptome energy scores (TES) between two phases in the life cycles of *B. bacteriovorus*. Calculations were performed (**a**) for all genes that are expressed per phase and (**b**) only for phase-specific genes. The TES value represents the average opportunity cost of a gene product multiplied by its frequency in the transcriptome at a given life cycle phase (Equation 3, Table A1). The significance of the difference between the two phases was tested by the Mann-Whitney U test with continuity correction. We depicted the corresponding W-value, p-value, and effect size (r). Outliers were removed from the graph for the clarity of presentation, but were included in the calculation of statistics. The X symbol represents the mean, and the horizontal line within a violin-plot represents the median. Transcription data was retrieved from Karunker et al. (2013) [14].

The attack phase is generally characterized by a much smaller number of expressed genes, which are more evenly distributed across the narrower range of TES values (Figure 5). In contrast, the TES values of the growth phase are predominantly grouped at the lower end of the TES range. This suggests that the attack phase of predatory *B. bacteriovorus* is underpinned by high transcription of a relatively small number of genes many of which encode expensive proteins. In contrast, the growth phase is characterized by relatively low transcription of many genes that encode cheaper proteins. These differences are even more apparent in the analysis of phase-specific genes (Figure 5b). Together, this suggests that the active predation in *B. bacteriovorus* requires proteins which are composed of expensive AAs. As an analogy, we previously speculated that similar phenotypes might exist in the context of animal predation [1].

### 2.6. Expensive Proteins Are Functionally Understudied

To investigate how Bdellovibrionota and Myxococcota differ in terms of the functions of their most expensive proteins, we conducted an enrichment analysis of COG functions (Figure 6). For each group, we separately calculated the opportunity cost (high respiration) of each protein (Equation 4) and used the MMseqs2 clustering approach [21] to group them into homologous clusters [5,8]. The opportunity cost of each cluster was then determined by averaging the opportunity costs of its members (Equation 5). Finally, we performed an enrichment analysis of COG functions for the top 10%, 20%, 30%, 40%, and 50% most expensive clusters within each clade (Figure 6).

**Figure 6.**
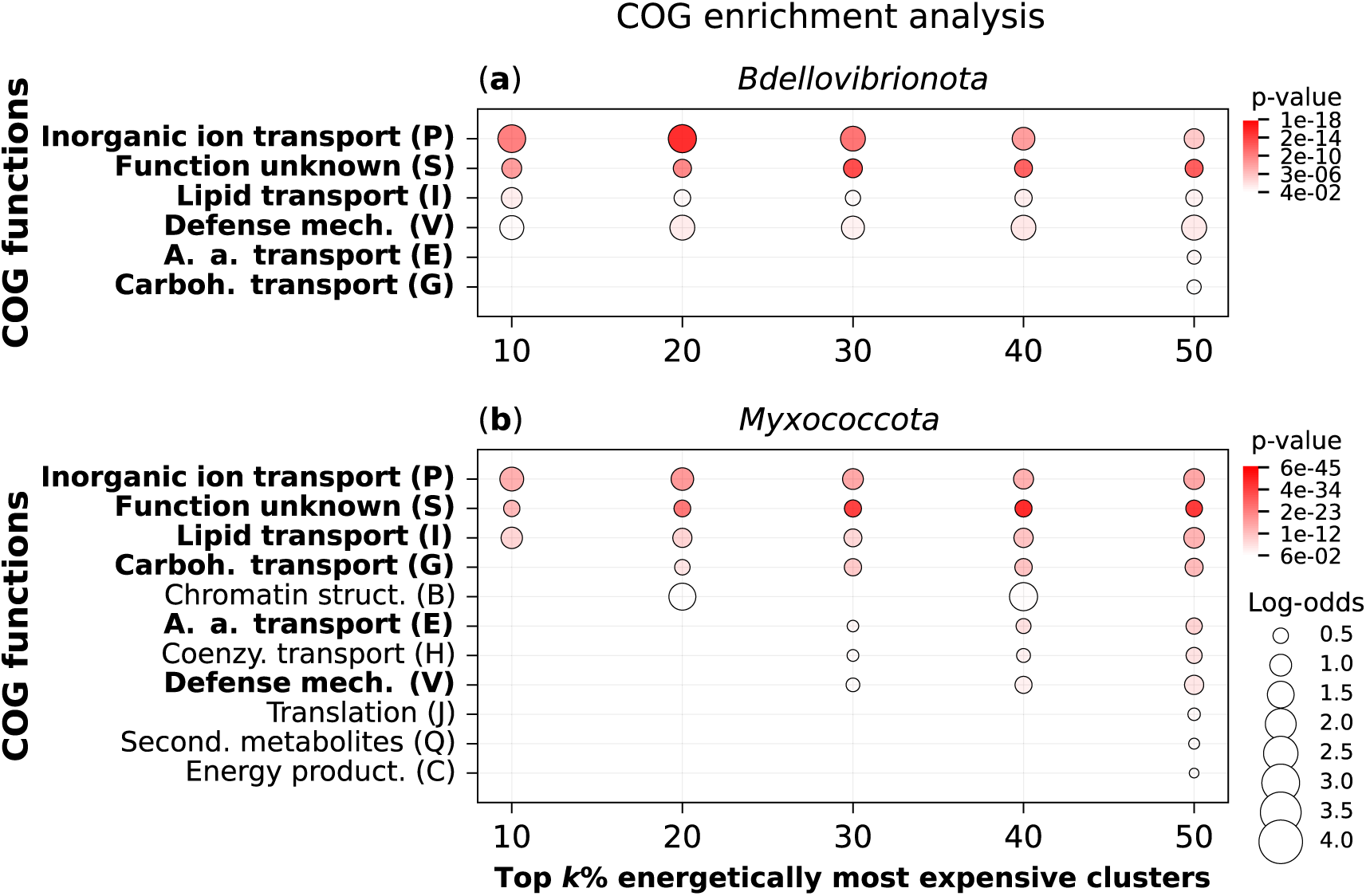
COG enrichment analysis of expensive protein clusters in Bdellovibrionota and Myxococcota. We analyzed the proteomes of (**a**) Bdellovibrionota (89 species), and (**b**) Myxococcota (203 species). Each dataset was clustered separately using the MMseqs2 algorithm to identify clusters of homologous proteins. COG functions were assigned to each cluster using EggNOG-mapper (see Materials and Methods). We performed an overrepresentation analysis for the top k% of clusters ranked by OC*_cluster_* (Equation 5, Table A1), with k = 10, 20, 30, 40, and 50, using a one-tailed hypergeometric test. The resulting *p*-values were corrected for multiple testing using the Benjamini–Hochberg method. Only enrichment signals with *p*-values < 0.05 are shown (Table S4).

The two groups show enrichment in a set of COG functions related to defense mechanisms, transport, and proteins of unknown function. These are the only functions enriched in Bdellovibrionota (Figure 6a), while in Myxococcota, we observe the enrichment of additional functions related to chromatin structure, translation, secondary metabolites, and energy production (Figure 6b). This suggests two key points: (i) a large number of expensive proteins in bacteria remain understudied, and (ii) the energetically demanding proteins encoded by Myxococcota are involved in a broader range of cellular functions compared to those of Bdellovibrionota.

## 3. Discussion

Functional outsourcing assumes that genes supporting essential functions can be lost from the genome if their activity can be substituted through environmental interactions [8]. We have successfully applied this concept to the evolution of AA auxotrophies in animals and bacteria [1,5]. We found that all animals and some bacterial groups which are capable of harvesting a sufficient amount of AAs from their ecosystem lost the ability to produce expensive AAs on their own [1,5,8]. This AA outsourcing is at least partially driven by energy-optimizing selection, which not only favors the loss of the ability to synthesize expensive AAs, but also allows for more frequent usage of expensive AAs in the proteome [1,5]. We proposed that in animals these phenotypes could have been triggered by predation and aerobic respiration [1]. If these processes are indeed selection-driven, it would be expected that they occur convergently under similar ecological pressures across the tree of life.

Unlike most other bacteria, Bdellovibrionota and Myxococcota are aerobic predators; they hunt and consume their bacterial prey under aerobic or microaerobic conditions [11–13,23]. As an independent evolutionary event, predation was likely crucial for the outsourcing of AA production in animals [1]. We have shown here that obligate aerobic predation left an astonishingly similar metabolic impact on Bdellovibrionota, resulting in the largely overlapping set of AA auxotrophies compared to animals. In contrast, very few auxotrophies can be observed in Myxococcota, which are facultative predators [18,20].

There are likely multiple factors that influenced the vast differences in AA biosynthesis capabilities between Bdellovibrionota and Myxococcota. However, the most apparent and potentially crucial factor is that Myxococcota are only facultatively predatory [18], preventing them from consistently obtaining AAs from the environment in sufficient quantities. This is supported by the observation that Bdellovibrionota grow and assimilate carbon at higher rates than Myxococcota [18]. Furthermore, it has been observed that Bdellovibrionota are significantly more abundant in aerobic environments than in anaerobic ones, in contrast to Myxococcota, which show no apparent preference [23]. This might indicate that the metabolism of Bdellovibrionota is more dependent on efficient respiration, which could also drive them toward increased AA auxotrophy levels [1]. Another important factor might be feeding efficiency—Bdellovibrionota are always physically connected to their prey, while Myxococcota secrete hydrolytic enzymes around their prey, which carries the risk of diffusion, leading to a lower return of the energy expended on predation [24].

The reduction in the ability to synthesize expensive AAs is directly correlated with the increased usage of expensive AAs in the proteomes [1,5]. Animals use more frequently expensive AAs in their proteomes than their sister group, choanoflagellates, and here we showed that the same is true for Bdellovibrionota when compared to their related group, Myxococcota [15]. Although Myxococcota have larger proteomes than Bdellovibrionota [11,24], their encoded AAs are on average cheaper by a large margin, which suggests that their complex lifestyle requires a wide range of functions that do not require proteins with very expensive AAs. This is supported by the results of our functional analysis, which showed that the top 50% most expensive proteins in Myxococcota are involved in a broader range of functions compared to the corresponding fraction of Bdellovibrionota proteins.

On the other hand, the ecology of Bdellovibrionota is relatively simple, consisting of two distinct phases: the attack phase, during which cells actively hunt, and the growth phase, during which cells consume their prey and subsequently divide [11,12,14]. A previous study produced transcriptome data for these two phases in *B. bacteriovorus* [14], providing a unique opportunity to examine the effects of energy-related selection on the *B. bacteriovorus* life cycle. We used a novel measure, the transcript energy score (TES), which combines the energy cost of a coding gene with its level of transcription, producing higher scores for more expensive and highly expressed transcripts. Using this metric, we found that the growth phase is underlined by a much broader set of transcripts with relatively similar expression levels that encode for cheaper AAs, while the attack phase consists of a relatively narrow subset of transcripts with a wide range of expression levels that encode for more expensive AAs. This supports the possibility that the energy saved through the outsourcing of AAs was invested in the attack phase, enabling more efficient predatory behavior and thus ensuring a more consistent influx of AAs from the environment, creating a positive feedback loop.

In conclusion, it is evident that every level of biological organization—from metabolism and proteome composition to the regulation of gene transcription—is intimately tied to the organism’s energy budget. Of course, other factors such as nutrient availability [5], ecological interactions [25], and horizontal gene transfer [26] can influence the pattern of AA auxotrophies, contributing to the complexity of metabolic evolution. However, the repeated convergent evolution of similar sets of AA auxotrophies across eukaryotes and prokaryotes, driven by obligate predation under aerobic conditions, suggests the impact of energy-related selection. This, in turn, appears to be linked with the evolution of novel and energy-expensive functions related to predation. Deeper understanding of the evolutionary pressures leading to predatory behavior in bacteria is especially important, as it has implications not only for understanding the regulation of ecological networks [18] but also for potential medical applications [27].

## 4. Materials and Methods

### 4.1. Databases, completeness score and auxotrophy index

All proteomes used in this study were retrieved from the NCBI GenBank. We acquired the highest-quality Bdellovibrionota (89) and Myxococcota (203) proteomes by using the NCBI filter to exclude atypical genomes, metagenome-assembled genomes, and genomes from large multi-isolate projects.

We conducted the detection of amino acid biosynthesis pathway completeness using the protocol described in our earlier publication [5]. We assessed the sensitivity of our method by comparing it to the results of in-vitro experiments. The error rate of our approach was comparable to other available in silico methods [5]. Briefly, we compiled a reference database of 387,892 enzyme sequences from 2,095 bacterial species, with each enzyme annotated according to the biosynthetic pathway it is involved in [5]. We combined the reference database with our proteomes and clustered the sequences using MMseqs2 [21] with the following parameters: –cluster-mode 0, –cov-mode 0, –c 0.8, and –e 0.001. We then functionally annotated all members of a cluster based on the presence of enzymes from the reference database in that cluster.

For each AA biosynthesis pathway and species in the database, we calculated a pathway completeness score (CS, Table A1) by dividing the number of detected enzymes by the total number of enzymes in that pathway, resulting in values ranging from 0 to 1. If a species contained alternative biosynthetic pathways for an AA, the pathway with the highest completeness score was selected. We then calculated the AA auxotrophy index of the *i*-th amino acid (*i =* 1, …, *20*) (*AI_i_*) by subtracting the completeness score (CS) from 1 [5]:

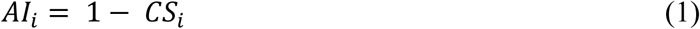

### 4.2. Opportunity cost measures, permutation and transcriptome analyses

We used the energy costs of AA biosynthesis as described in our earlier publications [1,5], details available in Table S3. The opportunity cost reflects the impact of AA synthesis on the cell’s energy budget and is calculated as the sum of the energy lost in the synthesis of AAs (direct cost) and the energy that would have been produced if a cell catabolized precursors instead of making AAs. Using the AA opportunity cost, we also calculated the opportunity cost of an average AA in each proteome (*OC_proteome_*) using the following equation:

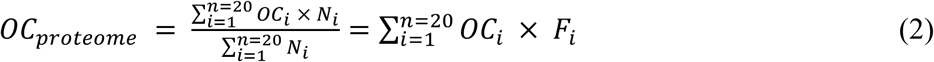

In this equation, OC*_i_* represents the opportunity cost of the *i*-th AA, *N_i_* denotes the total number of occurrences of this AA in the entire proteome, and *F_i_* represents the frequency of the AA in the proteome.

The permutation analyses were performed separately for each species by first determining the number of auxotrophic AAs and then generating all possible permutations for that number of auxotrophic AAs. For each permutation, we then calculated the average opportunity cost of the auxotrophic AAs. For example, if a species was found to be auxotrophic for 10 AAs, we found all possible combinations of 10 AAs and calculated the average opportunity cost of each. Since there is a limited number of possible average values, each value was treated as a bin. We calculated the proportion of permutations within a bin by dividing the number of elements in that bin by the total number of permutations. The obtained distribution represents empirical probability mass function (PMF) which was then used to calculate the probability that the observed set of auxotrophies in a given species is a result of a random process. We calculated p-values by summing the proportions of permutation in the range from the actual value observed in nature to the most extreme value at the closest distribution tail [1].

For transcriptome analysis, we obtained data on differential gene expression in the two distinct life cycle phases of *Bdellovibrio bacteriovorus* from an earlier study [14]. For each gene (*i*), we calculated its proportion in a given transcriptome phase by dividing its transcription value (*t_i_*) by the sum of all transcription values in that phase. This proportion in essence represents the frequency of each transcript in the transcriptome phase (*f_transcript(i)_*). We then multiplied this number by the opportunity cost (high respiration mode) of the protein encoded by that gene (OC*_protein(i)_*) to obtain a transcript energy score (TES). This score is meant to represent the energy impact of each transcript on the total energy budget of a transcriptome phase. TES of the *i*-th gene was calculated using the following equation:

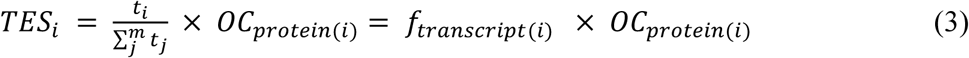

In this equation, *OC_protein(i)_* is the opportunity cost of the *i*-th protein (Equation 4), *f_transcript(i)_* is the transcription frequency of the *i*-th gene, *t_i_* denotes the transcription value of the *i*-th gene, *t_j_* denotes the transcription value of the *j*-th gene in the transcriptome, *j* = 1 to *m*, where *m* is the total number of transcribed genes.

The opportunity cost of proteins was calculated using the following equation:

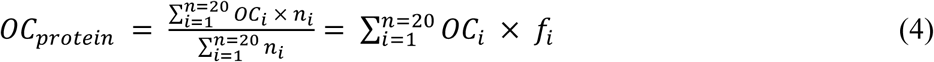

In this equation, *OC_protein_* is a weighted mean where *OC_i_* denotes the opportunity cost of the *i*-th amino acid, *n_i_* is the number of occurrences of the *i*-th of amino acid in a protein, and *f_i_* is the frequency of the *i*-th amino acid in a protein.

We used the Mann-Whitney U test with continuity correction to compare the opportunity costs of an average AA in proteomes (OC*_proteome_*) of Bdellovibrionota and Myxococcota using the package rcompanion (https://CRAN.R-project.org/package=rcompanion). We used the same test to compare transcript energy scores (TES) of different life cycle phases in *B. bacteriovorus*. To calculate correlations, we used the cor.test() function in the R stats (version 3.6.2) package. The heatmap was visualized using the ComplexHeatmap package [28].

### 4.3. COG functions enrichment analyses

We functionally analyzed the full proteomes of Bdellovibrionota (89) and Myxococcota (203). The two datasets were clustered separately using the MMseqs2 clustering algorithm (version 14-7e284) with the following parameters: –e 0.001 –c 0.8 –-max-seqs 400 –-cluster-mode 1 [8,21]. For each protein in the datasets, we obtained COG annotations using EggNOG-mapper (version 2.1.12) [29] with the DIAMOND (version 2.1.8) search tool [30]. We also calculated the opportunity cost (OC*_protein_*) for each protein under high respiration mode [1] (see Equation 4).

Finally, we performed the functional enrichment analysis on the two datasets independently. For each dataset, a cluster was assigned a COG function if at least one of its members was annotated with that function. The enrichment analysis was conducted for clusters with at least 10 members. For each cluster, we calculated the average opportunity cost (OC*_cluster_*) as follows:

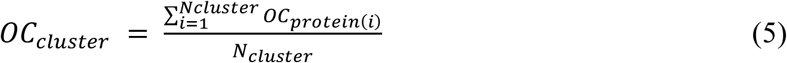

In this equation, *OC_protein(i)_* is the opportunity cost of the *i*-th protein in a cluster (see Equation 4) and *N_cluster_* is the number of proteins in the cluster.

We performed an overrepresentation analysis for the top k% of clusters ranked by OC*_cluster_*, with *k* = 10, 20, 30, 40, 50, using a one-tailed hypergeometric test as implemented in the Python (version 3.12.6) *scipy.stats* module. The obtained *p*-values were corrected for multiple testing and adjusted using the Benjamini–Hochberg method as implemented in the Python *statsmodels* library [31]. All results of enrichment analysis are shown in Table S4.

## Supplementary Materials

File S1: Full pathway completeness heatmap; File S2: Per-species statistics on binary encoded auxotrophies; Table S1: Database with pathway completeness score; Table S2: Summary statistics of binary auxotrophies; Table S3: Amino acid biosynthesis costs; Table S4: Enrichment analysis.

## Author Contributions

Conceptualization, N.K., M.D.-L. and T.D.-L.; methodology, N.K., M.D.-L. and T.D.-L.; software, N.K. and M.D.-L.; validation, N.K., M.D.-L. and T.D.-L.; formal analysis, N.K., M.D.-L. and T.D.-L.; writing—original draft preparation, N.K., M.D.-L. and T.D.-L.; writing—review and editing, N.K., M.D.-L. and T.D.-L.; visualization, N.K. and M.D.-L.; supervision, M.D.-L. and T.D.-L. All authors have read and agreed to the published version of the manuscript.

## Informed Consent Statement

Not applicable.

## Data Availability Statement

All data are available in the Supplementary Materials.

## Acknowledgments

We thank M. Futo, A. Tušar, S. Koska, D. Franjević and G. Klobučar for discussions. We used the computational resources of the University Computing Center – SRCE (Padobran) and the Institute Ruđer Bošković.

## Appendix A

**Table A1.** Abbreviations and acronyms.

| Abbreviation | Meaning | Equation |
| --- | --- | --- |
| AA | Amino acid | - |
| AAs | Amino acids | - |
| AI | Auxotrophy index | Eq. 1 |
| CS | Completeness score | Eq. 1 |
| $AI_i$ | Auxotrophy index of the $i$-th amino acid | Eq. 1 |
| OC | Opportunity cost | Eq. 2 |
| $OC_{protein}$ | Protein opportunity cost (weighted mean) | Eqs 3, 4, 5 |
| $OC_i$ | Opportunity cost of the $i$-th amino acid | Eqs 2, 4 |
| $n_i$ | The number of occurrences of the $i$-th amino acid in a protein | Eq. 4 |
| $f_i$ | The frequency of the $i$-th amino acid in a protein | Eq. 4 |
| $OC_{proteome}$ | Proteome opportunity cost (weighted mean) | Eq. 2 |
| $N_i$ | The number of occurrences of the $i$-th amino acid in a proteome | Eq. 2 |
| $F_i$ | The frequency of the $i$-th amino acid in a proteome | Eq. 2 |
| TES | Transcript energy score | Eq. 3 |
| $TES_i$ | The $i$-th gene transcript energy score | Eq. 3 |
| $t_i$ | Transcription value of the $i$-th gene | Eq. 3 |
| $f_{transcript(i)}$ | The transcription frequency of the $i$-th gene in a transcriptome | Eq. 3 |
| $OC_{cluster}$ | Cluster opportunity cost (average) | Eq. 5 |
| $N_{cluster}$ | Number of proteins in a cluster | Eq. 5 |

